# Early detection of active Human Cytomegalovirus infection after living-donor liver transplantation

**DOI:** 10.1101/304360

**Authors:** Mohamed Anies Rizk, Salah Abd Elfatah Agha, Maysaa El Sayed Zaki, Noha Badr El-Deen El-Mashad, Mohamed Mahmoud Fahmy El-Saadany

**Affiliations:** Clinical Microbiology Unit, Clinical Pathology Department, Faculty of Medicine, Mansoura University, Egypt.; Clinical Pathology Department, Faculty of Medicine, Mansoura University, Egypt.; Internal Medicine Department, Faculty of Medicine, Mansoura University, Egypt.

**Author notes:** Corresponding author e. mail.

**Keywords:** HCMV: Human Cytomegalovirus, LDLT: Living-donor liver transplantation, PCR: Polymerase Chain Reaction., ELISA: Enzyme linked Immunosorbent Assay

## Abstract

Human cytomegalovirus (HCMV) is a member of the beta herpes virinae. It is one of the most important virus in transplantation. It has direct and indirect impact on liver transplant recipient outcome. We aimed to diagnose early active HCMV infection in living donor liver transplant (LDLT) recipients. Also, to correlate the associated clinical and laboratory findings with HCMV infection. Here, we investigate 76 LDLT recipients for early detection of active HCMV infection in a period of 1-6 months after liver transplantation upon their suggested clinical data. These samples were collected in the period from 4/2013 to 12/ 2015 at Gastroenterology center, Mansoura University. They were 68 males and 8 females. HCMV infection diagnosed by ELISA (Ig M, Ig G) and real time PCR. Seventy-four patients were IgG seropositive recipient. Three patients (3.9%) were positive IgM. Ten samples from 76 patients (13.2%) were positive by real-time polymerase chain reaction (PCR). In this study, we concluded that, LDLT recipients are at high risk of HCMV infection (13.2%) and the most suitable method for HCMV detection was PCR.

## Introduction

Opportunistic viral infections are common after liver transplantation. The herpes virus infections are common after transplantation; Human cytomegalovirus (HCMV) is most important virus of this group ^(1)^. HCMV belongs to the beta herpes virus family. Beta herpes viruses are closely related to each other with large genomic overlapping ^(2)^.

The great seroprevalence and early acquisition of the virus have been associated with lower socioeconomic circumstances, developing countries and over crowded populations ^(3)^. HCMV infects 60%-100% of humans, with primary HCMV infection occurring most commonly during the first two decades of life. In immunocompetent individuals, the infected individuals are mostly asymptomatic or may present with a benign febrile infectious mononucleosis-like illness ^(4)^. HCMV infection occur in the majority of solid organ transplant patients, primarily in the first months when immunosuppression is most intense, HCMV disease incidence ranges from 10% to 50% ^(4)^.

Liver transplant recipients are at a higher risk of HCMV infection, especially those who did not have HCMV infection until they receive a latently infected organ from a HCMV-seropositive donor (HCMV D+/R-mismatch). The risk of progression into HCMV disease is magnified by the intense immunosuppression required to avoid or to treat allograft rejection ^(5)^. The clinical impact of HCMV disease after liver transplantation can be classified into: (1) an acute infection with clinical signs known as direct effects (fever, mononucleosis, and invasive organ disease); and (2) a broad range of immunomodulatory and vascular effects, referred to as indirect effects ^(6)^.

The aim of this study was to detect the incidence of HCMV infections in living-donor liver transplantation recipients by ELISA and real time PCR. Also, to correlate the laboratory findings with HCMV infections.

## Materials and Methods

### Subjects

This study was conducted on 76 living-donor liver transplant (LDLT) recipients admitted to the Gastroenterology Surgery Center, Mansoura University in the period from April 2013 to December 2015. They were 68 males and 8 females. Written consent was obtained from patients. They were investigated for detection HCMV infections in a period up to 6 months after liver transplantation according to suspected clinical symptoms (e.g. fever, hepatitis, pneumonitis, urinary tract infections, gastroenteritis such as diarrhea and vomiting) (**Group I**). In addition, 10 living-donor liver transplant recipients of matched age and sex, who did not complain of any symptoms suggestive of HCMV, were included as a control group (**Group II**).

### Methods

**I.For the patients and control group, the followings were done:**

- History was taken, thorough clinical examination, to detect suspected cases, e.g. (fever, gastroenteritis as; diarrhea, vomiting, urinary tract infections, pneumonitis).
- Routine laboratory investigations as; Liver function tests (Bilirubin, Albumin, ALT, AST, ALP and GGT), kidney function tests (creatinine, urea and uric acid), C-reactive protein (CRP) were carried out using (Cobas Integra 800, Roche Diagnostics), and complete blood picture (CBC) was carried out using (Sysmex 1000).

**II. Detection of HCMV antibodies (IgM, IgG) by ELISA**

**II. a. Detection of HCMV IgM and IgG:** (DRG International, Inc., USA)

Blood samples were collected in a sterile vaccutainers and the serum was separated and stored at −20 °C until processing.

**II. b. Molecular detection of HCMV DNA by real-time PCR:**

Blood samples were collected in a sterile vaccutainers and the serum was separated and stored at −20 °C until processing.

**II. b. 1. DNA extraction:**

HCMV DNA was extracted using QIAamp mini kit (Cat. No. 57704 QIAamp miniElute virus spin kit, QIAGEN GmbH, QIAGEN Strasse 1, D-40724 Hilden, Germany).

**II. b. 2. HCMV DNA amplification and detection by real-time PCR.**

The Artus CMV TM PCR Kit (QIAGEN GmbH, QIAGEN Strasse 1, D-40724 Hilden, Germany) was used for detection of CMV-DNA.

#### Principle

HCMV detection by the polymerase chain reaction (PCR) is based on the amplification of specific regions of HCMV genome. In real-time PCR, the amplified product is detected via fluorescent dyes. These are usually linked to oligonucleotide probes, which bind specifically to the amplified product.

#### Amplification and detection

Amplification and detection performed using the STRATAGENE (Applied Biosystems Inc., Foster City, CA)

#### Thermal profile

Initially precycling denaturation and then Taq activation achieved at 95 °C for 10 minutes. Then 45 cycles of denaturation at 95 °C for 15 seconds, annealing and extension at 55 °C for 1 minute (Table I).

**Table I:**
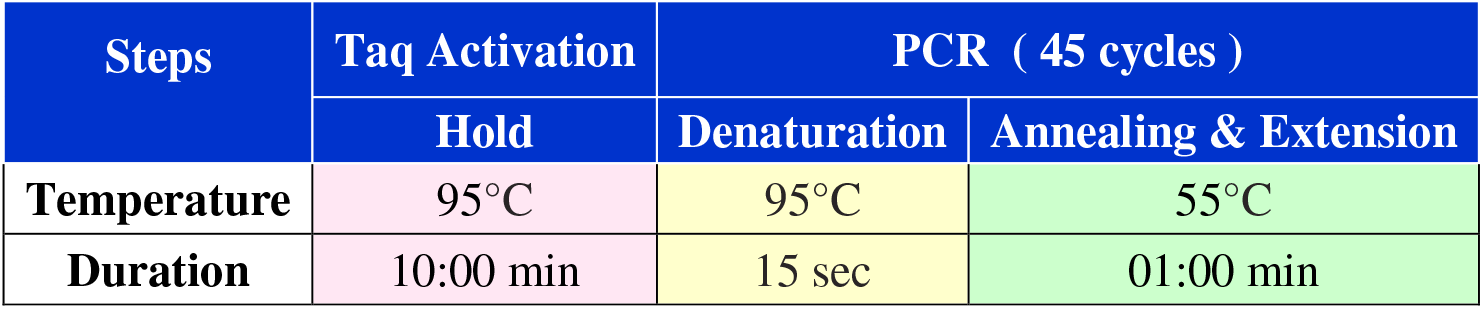
Thermal profile of HCMV DNA real-time PCR (Applied Biosystems Inc., Foster City, CA)

The results were then quantified using HCMV DNA at predetermined concentrations provided by the manufacturer or from standard curve (Figure I).

**Figure I:**
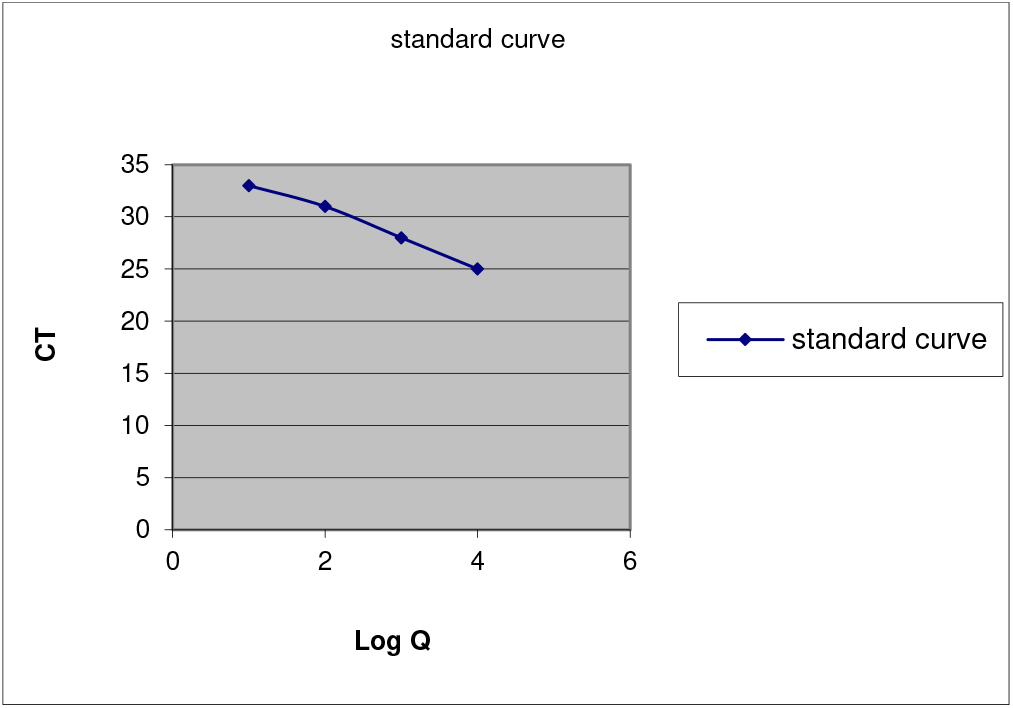
Standard curve of quantitation of HCMV by real-time PCR

## Statistical analysis

Data analyzed and processed with SPSS software (Statistical Product and Service Solutions, version 18.0, SSPS Inc, Chicago, IL, USA). Qualitative data expressed in frequency and percentage and analyzed with the chi-square test or the Fisher exact test, when appropriate. Quantitative data expressed as the mean and standard deviation and compared with the t test. The correlation between two quantitative variables done with Pearson’s correlation coefficient or Spearman’s rank correlation coefficient, when appropriate. In all tests, a P value of < 0.05 was considered significant, and a P value of < .01 was highly significant.

## Results

The age and sex distribution in the studied patients ranged from 29 to 63 years old with a mean of 51.14±7.116. They were 68 males and 8 females. The mean age of the control group (group II) was (49.90±6.74) (Table 2). The mean time for occurrence of active HCMV infections occurred during the second post-transplant month (2.58±5.347) (Table 3).

**Table (2):** Age and sex distribution in all studied subjects.

|  |  | **Group I**<br>**(n=76)** | **Group II**<br>**(n=10)** | ***p*** |
| --- | --- | --- | --- | --- |
| **Age (years)** |  | 51.14±7.116 | 49.90±6.740 | 0.602 |
| **Sex** | **Males** | 68 (89.5%) | 8 (80.0%) | 0.328 |
|  | **Females** | 8 (10.5%) | 2 (20.0%) |  |

P: significance between total cases versus control.

**Table (3):** Time-related HCMV infection in living-donor liver transplanted patients.

|  | **Group I**<br>**(n=76)** | **Group II**<br>**(n=10)** |
| --- | --- | --- |
| **HCMV** | 2.58±5.347 | 1.6±0.699 |

The clinical data of the studied patients. Fever was the most common presentation in all studied cases (69.7%) followed by hepatitis (56.6%), gastroenteritis (42.1%), pneumonitis (5.3%). There was a significant difference between patients versus control group regarding to gastroenteritis, hepatitis and fever (p=0.011, 0.001 and <0.001 respectively) (Table 4).

**Table (4):**
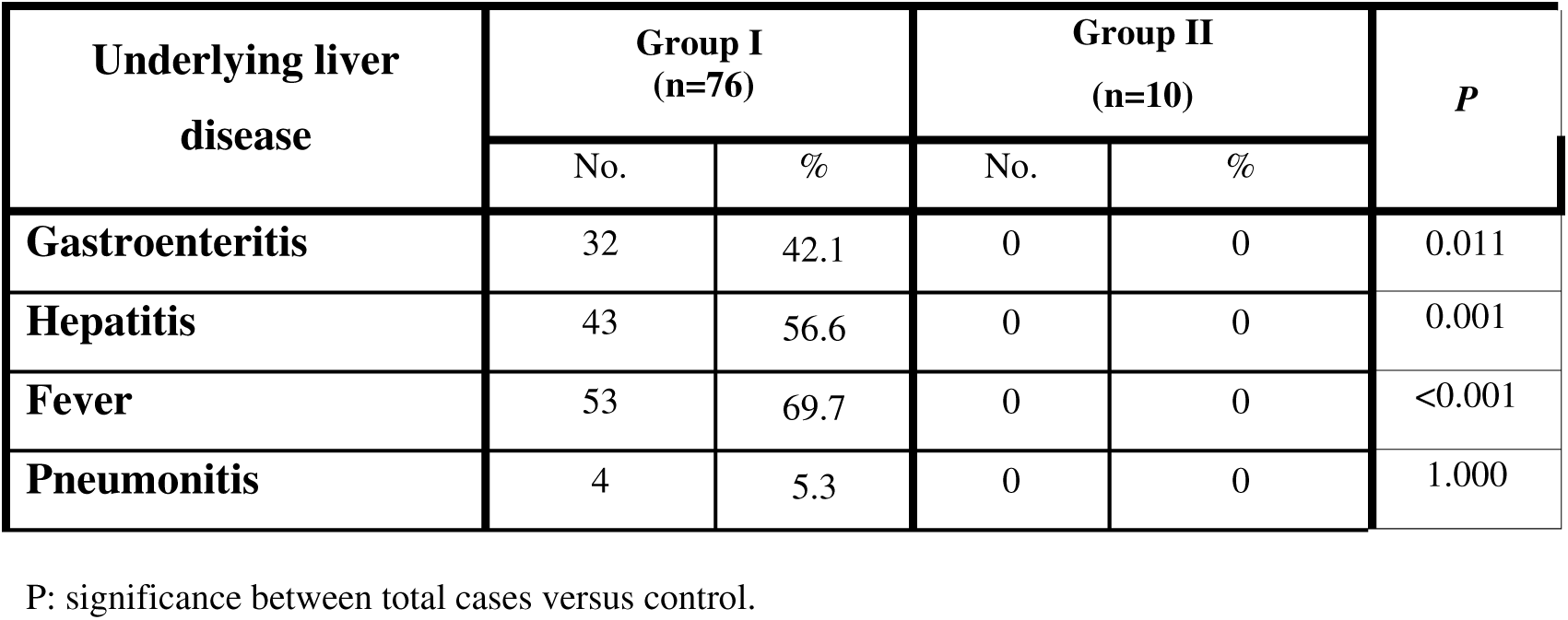
Clinical data of the studied patients.

| Underlying liver disease | Group I (n=76) |  | Group II (n=10) |  | *P* |
| --- | --- | --- | --- | --- | --- |
|  | No. | % | No. | % |  |
| **Gastroenteritis** | 32 | 42.1 | 0 | 0 | 0.011 |
| **Hepatitis** | 43 | 56.6 | 0 | 0 | 0.001 |
| **Fever** | 53 | 69.7 | 0 | 0 | <0.001 |
| **Pneumonitis** | 4 | 5.3 | 0 | 0 | 1.000 |

P: significance between total cases versus control.

HCMV detection by different methods in the studied cases: out of 76 patients, 74 (97.45 %) were IgG seropositive, 3 (3.9%) were positive for IgM and 10 (13.2%) were positive by real-time PCR. They were compared to control subjects and there was no significant difference as regard IgG. On the other hand, there was significant difference as regards IgM and PCR (Table 5).

**Table (5):** HCMV detection rate by different methods in studied cases (n=76).

|  |  | Group I (n=76) |  | Group II (n=10) |  | *P* |
| --- | --- | --- | --- | --- | --- | --- |
|  |  | No | % | No | % |  |
| **CMV IgG (ELISA)** | **Positive** | 74 | 97.4 | 10 | 100.0 | 1.000 |
|  | **Negative** | 2 | 2.6 | 0 | 0 |  |
| **CMV IgM (ELISA)** | **Positive** | 3 | 3.9 | 0 | 0 | 1.000 |
|  | **Negative** | 73 | 96.1 | 10 | 100.0 |  |
| **CMV DNA Real time PCR** | **Positive** | 10 | 13.2 | 0 | 0 | 0.588 |
|  | **Negative** | 66 | 86.8 | 10 | 100.0 |  |

P: significance between total cases versus control.

The sensitivity, specificity, positive and negative predictive values of ELISA IgM for HCMV in comparison to real-time PCR as the reference standard. Both techniques gave positive results in two cases and negative in 65 patients. The sensitivity, specificity, positive predictive value and negative predictive value of ELISA IgM for HCMV were (20.0%, 98.5%, 66.7% and 89.0% respectively). Prevalence of positive HCMV in studied group I= 13.2% (Table 6).

**Table (6):** Sensitivity, specificity, positive and negative predictive values of ELISA IgM for detection of HCMV in comparison PCR.

| Method of Detection |  | Negative HCMV (66) |  | Positive HCMV (10) |  | Sensitivity (%) | Specificity (%) | PPV (%) | NPV (%) |
| --- | --- | --- | --- | --- | --- | --- | --- | --- | --- |
|  |  | No | % | No | % |  |  |  |  |
| CMV IgM (ELISA) | Positive (n=3) | 1 | 1.5 | 2 | 20.0 | 20.0 | 98.5 | 66.7 | 89.0 |
|  | Negative (n=73) | 65 | 98.5 | 8 | 80.0 |  |  |  |  |

PPV: Positive predictive value, NPV: Negative predictive value.

Clinical data of HCMV in studied group I (n=76). Gastroenteritis, hepatitis and fever showed significant differences between positive cases versus control subjects (p=0.033, 0.003, 0.001 respectively). Pneumonitis did not differ significantly between positive cases and control subjects (Table 7).

**Table (7):** Clinical data of HCMV infection in-group I (n=76).

| Clinical presentations | Negative HCMV (66) |  | Positive HCMV (10) |  | Group II (n=10) |  | P |
| --- | --- | --- | --- | --- | --- | --- | --- |
|  | No. | % | No. | % | No. | % |  |
| Gastroenteritis | 27 | 40.9 | 5 | 50.0 | 0 | 0 | 0.033 |
| Hepatitis | 36 | 54.5 | 7 | 70.0 | 0 | 0 | 0.003 |
| Fever | 45 | 68.2 | 8 | 80.0 | 0 | 0 | 0.001 |
| Pneumonitis | 2 | 3.0 | 2 | 20.0 | 0 | 0 | 0.474 |

P: significance between total cases versus control.

Clinical indications did not show significant differences between patients with positive HCMV infection versus those with negative HCMV infection and versus control subjects (p>0.05) (Table 8).

**Table (8):**
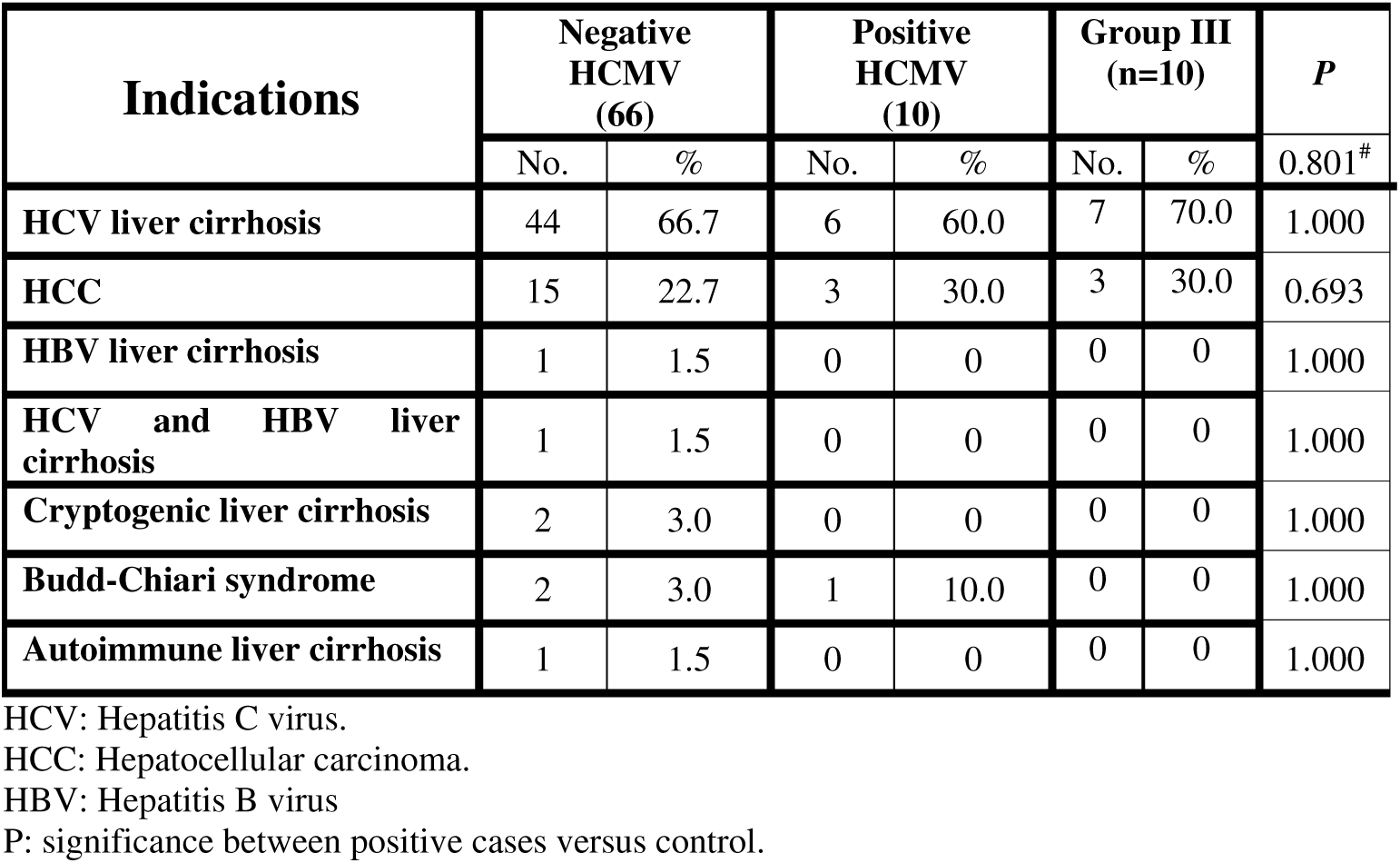
Relationship between clinical indications for liver transplantation to HCMV infection in all patients (n=76) versus control (n=10).

The relation between HCMV infection and laboratory data was increased serum bilirubin (total and direct) and GGT showed high significance differences between group I and control group (group II) (p=0.002, 0.001, 0.009 respectively), increased ALP and glucose showed marginally significant differences (p=0.093, 0.071 respectively), increased CRP showed a significant difference (p=0.038). Mild reduction of hemoglobin concentration and mild elevation of INR showed marginally significance (p=0.0503, 0.066 respectively) (Table 9).

**Table (9):** Comparison between HCMV infection and other laboratory data in group I (n=76) versus control (group II: n=10).

| Laboratory data |  | Negative HCMV<br>(n=66) |  | Positive HCMV<br>(n=10) |  | Group II<br>(n=10) |  | *P* | *P\** |
| --- | --- | --- | --- | --- | --- | --- | --- | --- | --- |
|  |  | Mean | SD | Mean | SD | Mean | SD |  |  |
| Albumin (g/dl) |  | 3.55 | 0.49 | 3.66 | 0.62 | 3.83 | 0.440 | 0.512 | 0.490 |
| Bilirubin | Total (mg/dl) | 2.93 | 5.92 | 2.19 | 1.02 | 0.99 | 0.179 | 0.697 | 0.002 |
|  | Direct (mg/dl) | 1.85 | 4.91 | 1.31 | 0.80 | 0.15 | 0.053 | 0.729 | 0.001 |
| ALT (U/ml) |  | 50.26 | 94.79 | 28.20 | 11.72 | 26.90 | 6.707 | 0.467 | 0.764 |
| AST (U/ml) |  | 78.21 | 149.81 | 34.80 | 13.43 | 29.30 | 8.042 | 0.365 | 0.281 |
| GGT (U/L) |  | 152.48 | 138.61 | 111.30 | 81.38 | 25.20 | 5.266 | 0.364 | 0.009 |
| ALP (KAU) |  | 11.33 | 9.18 | 8.70 | 2.54 | 11.50 | 4.301 | 0.373 | 0.093 |
| Creatinine (mg / dl) |  | 1.24 | 0.69 | 1.06 | 0.62 | 0.91 | 0.145 | 0.450 | 0.464 |
| F.B.Glucose (mg / dl) |  | 138.83 | 61.09 | 140.00 | 64.52 | 100.70 | 6.056 | 0.956 | 0.071 |
| CRP (mg / L) |  | 41.95 | 53.92 | 38.40 | 45.56 | 3.50 | 1.080 | 0.844 | 0.038 |
| TLC (X10<sup>9</sup>/l) |  | 8.25 | 7.07 | 6.60 | 3.15 | 6.740 | 2.202 | 0.474 | 0.912 |
| Hb (g/dl) |  | 10.29 | 2.02 | 10.13 | 1.13 | 11.270 | 1.295 | 0.720 | 0.0503 |
| Platelets (X10<sup>9</sup>/l) |  | 157.50 | 88.48 | 204.80 | 116.24 | 165.00 | 68.323 | 0.135 | 0.363 |
| INR |  | 1.28 | 0.63 | 1.31 | 0.38 | 1.060 | 0.070 | 0.897 | 0.066 |

P: significance between total cases versus control.
P\*: significance between positive cases versus control.

The relationship of post-transplant complications with active HCMV in-group I (n=76) versus control group. Recurrent HCV had higher incidence of HCMV infection (40.0% versus 12.1% in positive versus negative HCMV, p=0.046). Cases complicated with acute rejection had higher incidence of HCMV infection (50.0% versus 24.2% % in positive versus negative HCMV), (p=0.128). No significant differences were found in recurrent HCC, biliary complications, other infections and pancytopenia (Table 10).

**Table 10.**
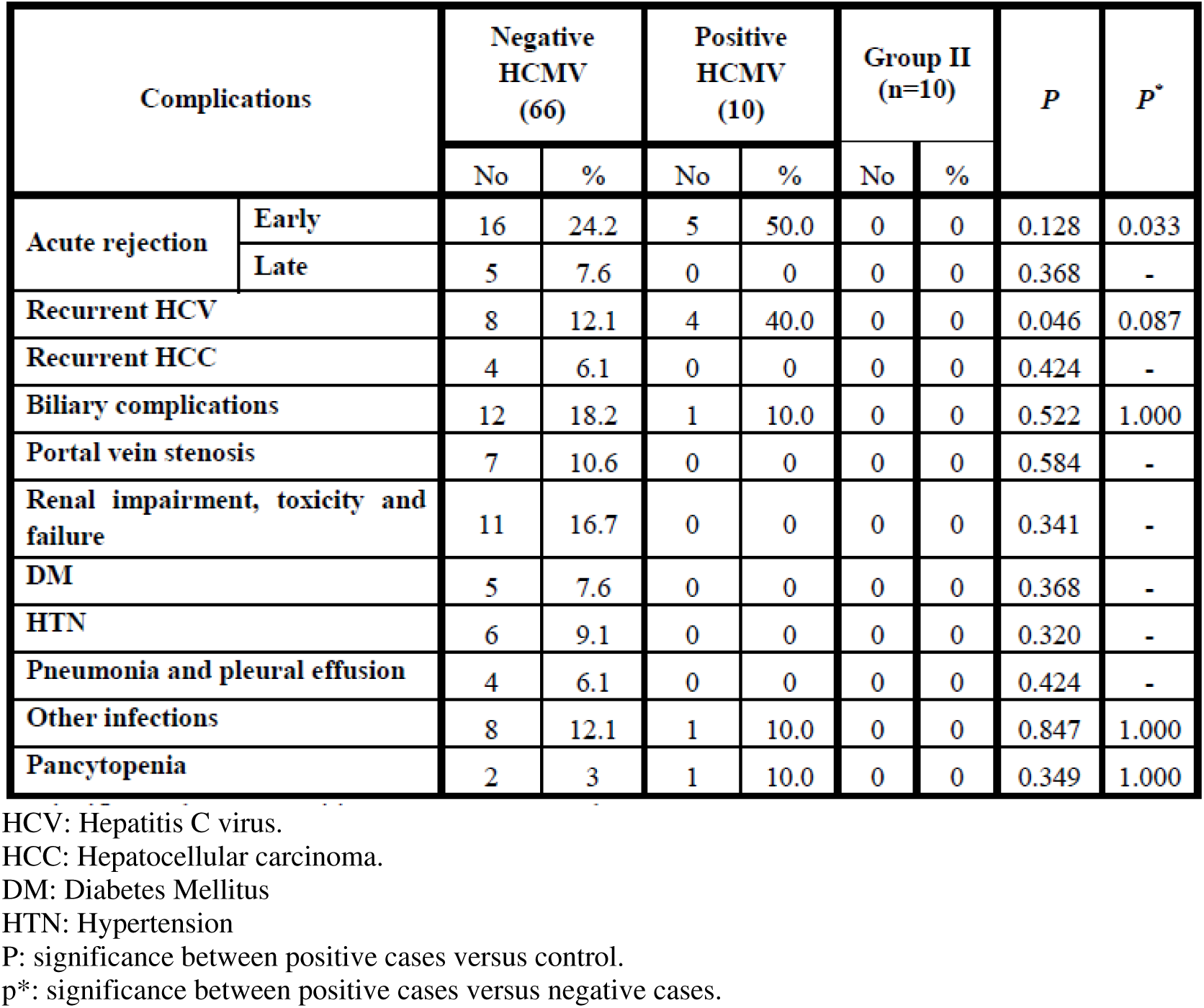
:Relationship of post-transplant complications to active HCMV in group I (n=76).

## Discussion

The age of the studied patients ranged from 29 to 63 years with a mean 51.14±7.116 years. They were 68 (89.47 %) males and 8 (10.52%) females. We investigate 76 patients for HCMV antibodies (IgM and IgG) and real-time PCR assays. Out of 76 patients, 74 (97.36 %) patients were IgG seropositive and three patients (3.9%) were HCMV IgM seropositive. Real-time PCR was positive in 10 out of 76 patients (13.2%). The high prevalence of HCMV IgG antibodies in this study indicates that the incidence of HCMV infection is high in Egypt and this may be attributed to the poor socioeconomic status and hygienic practices.

***Razonable and Pya, (2003)*** showed in their study that the incidence of HCMV infection ranges from 10% to 50% following solid organ transplantation ^(4)^. ***Jain et al., (2005)*** showed that, the incidence of HCMV infection was 14% after liver transplantation ^(7)^. In accordance ***Wadhawan et al., (2012)*** reported that HCMV reactivation was in 13% after LDLT, also, ***Katsolis et al., (2013)*** showed that HCMV infection was in 40% and disease was in 19% ^(8,9)^. In a study by ***Garbinoa et al. (2005)***, they reported that infection episodes among the 78 out of 98 patients (80%), 44% were due to viral infections and HCMV infection was present in 29% of these viral infections ^(10)^. ***Milan et al., (2013)*** documented that 16 (32%) patients had HCMV infections after liver transplantation, including 13 (81%) with concomitant infections. Thirty-four patients (68%) did not have HCMV infections and eight of them (24%) had bacterial infection. There was a high correlation with bacterial infections among CMV-positive patients ^(11)^.

We found two cases were positive by HCMV IgM and real-time PCR and 65 were negative by both techniques. The sensitivity, specificity, positive predictive value and negative predictive value of ELISA IgM for HCMV were (20.0%, 98.5%, 66.7% and 89.0%) respectively. The incidence of active HCMV infection by real-time PCR in studied group I= 13.2%.

***Martín-Dávila et al.(2004)*** showed that The sensitivity, specificity, positive predictive value (PPV) and negative predictive value (NPV) of real-time PCR were 78%, 77%, 47%, 89% respectively ^(12)^. The study by ***Schmidt et al., (1995)*** for diagnosis of HCMV infection after liver transplantation, showed that in a total of 30 patients, fourteen patients (46%) were positive by PCR. The sensitivity and specificity of PCR was 100% and 76%, respectively. However, for the HCMV IgM, the sensitivity was only 66%, and the specificity was 84%.The demonstration of HCMV IgM antibodies is of little practical help as the antibody response develops too late in relation to infection ^(13)^. ***Patel et al., (1994)*** reported that the sensitivity of PCR in the diagnosis of symptomatic HCMV infection in liver transplant recipients was 100%, the specificity was 57% and the positive predictive value was 50% ^(14)^.

In this study, there was no significant differences between patients with positive HCMV infection versus those with negative HCMV infection as regards clinical data (p>0.05). Gastroenteritis (50%), hepatitis (70%) and fever (80%) showed significant differences between positive cases versus control subjects (p=0.033, 0.003, 0.001 respectively). Pneumonitis (20%) did not differ significantly between positive cases and control subjects. The classic illness caused by HCMV after transplantation is manifested by fever, bone marrow suppression, and Invasive diseases, which had been traditionally categorized either as HCMV syndrome. The most common organ system involved during HCMV disease is the gastrointestinal tract (in the form of HCMV gastritis, esophagitis, enteritis, and colitis), accounting for over 70% of tissue invasive HCMV disease cases in solid organ transplant recipients ^(15)^. The clinical indications for liver transplantation did not show any significant differences between patients with positive HCMV infection versus those with negative infection or with control group (p>0.05).

The similarity of clinical indications for liver transplantation between the studied patients and control group might explain the insignificant difference between these groups. The incidence of recurrent HCV was higher in patients with active HCMV infection versus control 40.0% and 12.1% in negative HCMV (p=0.046) respectively. In addition, patients with active HCMV infection showed higher incidence of acute rejection compared to control group (50.0% and 24.2% %) respectively. On contrary, ***Singh et al., (2005)*** did not find an association between recurrent HCV and HCMV infection after liver transplantation. ***Rosen et al. (1997)*** reported that the incidence of HCMV induced cirrhosis was 50% compared to 11% in non-infected patients ^(16, 17)^. HCMV infection has been associated with an increased risk for severity of HCV recurrence and this might be related to immunosuppression induced by HCMV infection or immunosuppressive drugs ^(18)^. ***Singh et al., (2005)*** showed that, recurrent HCV infection after LDLT occurred in 55.6% of the patients with HCMV infection and 49.8% of patients without HCMV infection ^(16)^.

The HCMV positive cases in comparison with control group showed elevated serum bilirubin (total and direct) and serum GGT (p=0.002, 0.001, 0.009) respectively, and elevated serum ALP and hyperglycemia (p=0.093, 0.071) respectively. In addition, elevated level of CRP showed significant differences (p=0.038). Mild decrease in hemoglobin concentration and mild elevation of INR showed mild significant difference (p=0.0503, 0.066) respectively. In multivariable analysis, increased risk for HCMV infection was evident in patients with lower model for end-stage liver disease (MELD) score (P = 0.025), lower total bilirubin (P = 0.014), and longer operative time (P = 0.038) ^(9)^.

## Summary and conclusion

Out of 76 patients, 74 (97.36 %) patients were IgG seropositive and three patients (3.9%) were HCMV IgM seropositive. Real-time PCR was positive in 10 patients (13.2%). In this study, two cases were positive by HCMV IgM and real-time PCR and 65 were negative by both techniques. The sensitivity, specificity, positive predictive value and negative predictive value of ELISA IgM for HCMV were (20.0%, 98.5%, 66.7% and 89.0%) respectively. The incidence of active HCMV by real-time PCR in studied group I= 13.2%. Gastroenteritis (50%), hepatitis (70%) and fever (80%) showed significant differences between positive cases versus control subjects (p=0.033, 0.003, 0.001 respectively). The incidence of recurrent HCV was higher in patients with active HCMV infection versus control 40.0% and 12.1% in negative HCMV (p=0.046) respectively. In addition, patients with active HCMV infection showed higher incidence of acute rejection compared to control group (50.0% and 24.2% %) respectively. HCMV positive cases in comparison with control group showed elevated serum bilirubin (total and direct) and serum GGT (p=0.002, 0.001, 0.009) respectively, and elevated serum ALP and hyperglycemia (p=0.093, 0.071) respectively. According to laboratory investigations, elevated CRP showed significant differences (p=0.038). Mild decrease in hemoglobin concentration and mild elevation of INR showed mild significant difference (p=0.0503, 0.066) respectively.

